# How cancer patients perceive clinical trials (CTs) in the era of CTs: *Current perceptional status and differences between common and rare cancers*

**DOI:** 10.1101/455899

**Authors:** Ji Hyun Park, Ji Sung Lee, HaYeong Koo, Jeong Eun Kim, Jin-Hee Ahn, Min-Hee Ryu, Sook-ryun Park, Shin-kyo Yoon, Jae Cheol Lee, Yong-Sang Hong, Sun Young Kim, Kyo-Pyo Kim, Chang-Hoon Yoo, Jung Yong Hong, Jae Lyun Lee, Kyung Hae Jung, Baek-Yeol Rhyoo, Tae Won Kim

**Author notes:** **Correspondence**: Tae Won Kim, M.D., Ph.D., Address: Department of Oncology, Asan Medical Center, University of Ulsan College of Medicine, 88 Olympic-ro 43-gil, Songpa-gu, Seoul 05505, Korea.

## Abstract

**Background:** As perception have been recently highlighted as critical determinants of clinical trials (CTs) in cancer patients, we evaluated current perceptional status of CTs in cancer patients, focusing on differences between common and rare cancers.

**Materials & Methods:** From November 2015 to May 2017, we prospectively surveyed patients who have received anti-cancer treatment at Asan Medical Center using the PARTAKE questionnaires.

**Results:** Among 333 respondents, 70.9% and 29.1% had common and rare cancers, respectively. While 87.7% and 75.3% of patients answered that they heard of and knew about CTs, willingness to participate in CTs was expressed only in approximately 56% of patients although willingness was significantly correlated with awareness and perception. Surprisingly, patients with rare cancers showed significantly lower levels of awareness and perception (64.2% vs 79.9%, p=0.003 and 77.3% vs 91.9%, p<0.001), and consequently less willingness (47.4% vs 58.9%, p=0.06) compared to patients with common cancers. In addition, cancer patients still harbored fear with concerns about safety and reward, and substantial ignorance and mistrust about voluntariness of CTs, which was more predominant in patients with rare cancers.

**Conclusions:** Present study identified relatively less willingness of CTs in cancer patients compared with generally favorable perception, and highlighted relative perceptional poverty in patients with rare cancers than those with common cancers. Further education and encouragement by research and public entities seem essential to raise motivation of CTs in cancer patients beyond good perception, especially for the patients with rare cancers.

## Introduction

Clinical trials (CTs) have decisively contributed to new drug development for cancer patients in modern oncology, catalyzing therapeutic advances in the last decade(1). In the era of CTs, perceptional parameters have been highlighted as a crucial determinant of CTs accrual based on a compelling relationship between awareness and subsequent willingness or participation in CTs(1-5). Interestingly, actual accrual rates were still disappointing in cancer patients although they mostly harbored generally positive view of CT(1, 6). These findings consistently suggested significance of collaborative education by research or public entities to raise patients’ perception and motivation of CTs(1, 5, 7).

Meanwhile, patients with rare cancers have been frequently excluded from conventional types of CTs due to their inherent barriers of small populations and minimal evidences of standardized treatments(8-10). Necessarily, they have suffered from relative therapeutic deprivation compared with patients with common cancers, more obviously in case of young and adolescent patients(11-15). However, with development of newer molecular targets and multi-omics, CTs have evolved into more intuitive and comprehensive forms(16), and they particularly benefited patients with rare cancers by potentiating opportunities of enrollments(10, 17-20). In addition, several scientific and administrative initiatives recently launched their international collaborations to mitigate organizational hurdles of CTs in rare cancers (17, 18, 21). Regardless of newly fostered milieu of CTs in rare cancers, perceptional aspects of CTs in patients facing rare cancers have been rarely explored thus far.(12, 14, 15).

In the present study, therefore, we investigated current perception of CTs and their significant determinants in Korean cancer patients using the validated Public Awareness of Research for Therapeutic Advancements through Knowledge and Empowerment (PARTAKE) survey, and focused on differences between patients with rare and common cancers.

## Material and methods

### Patients collection

From November 2015 to May 2017, we prospectively surveyed cancer patients who received anti-cancer treatment at Asan Medical Center (Korea, Seoul). Patients were enrolled regardless of their stage, type, or treatment, with an attempt to include all of the patients with rare tumors. All common and rare cancers were defined using the classification of the RARECARE study group(22). Investigators initially explained the purpose and scope of the study either at the outpatient clinic or ward, and a well-trained clinical research assistant (CRA) administered the survey to patients who agreed to participate, after obtaining written informed consent. The questionnaires were collected by the CRA after completion by each enrolled patient at the time of the patient’s next visit or before their discharge. Study-related medical records of enrolled patients were reviewed by an investigator for further analysis. The study protocol was reviewed and approved by the Institutional Review Board (IRB approval number: 2014-1061) of Asan Medical Center and was conducted in full accordance with the guidelines for Good Clinical Practice and the Declaration of Helsinki.

### PARTAKE survey

The PARTAKE survey was devised to evaluate the status and structures of awareness (heard of) and perception (know about) of clinical research. It is composed of 40 multiple-choice and open-ended questions, classified in four different categories for assessing individuals’ perceptions including trust in research entities, conduct of research, motivations for participation, and the value of clinical research. As described in our last survey study using PARTAKE(5), it was extensively validated after its development by various methods including literature review and consultation with relevant experts. Additionally, we performed our own validation process before its virtual administration in December 2014, which included translation and then back-translation with the provision of additional written instructions for administering the questionnaire and minimizing selection bias. Other details of our methodology are as described in our last study(5).

## Statistical analysis

Descriptive statistics were used to demonstrate the patients’ characteristics and to assess general outcomes of perception and participation in CTs. The chi-square or Fisher’s exact test was applied to identify influential determinants of perceptional parameters including awareness, perception and willingness to participate in CTs among patients’ clinical and demographic characteristics. Comparative analysis of awareness and perception between patients with common (common cancer group) and rare cancers (rare cancer group) was performed with the chi-square or Fisher’s exact test based on the PARTAKE questionnaire. Additionally, perceptional outcomes of cancer patients were compared with general Korean public for a supplement. All statistical analyses were executed with SAS software (version 9.4, SAS Institute, Cary, NC, USA), and were performed by an independent statistician at our institute.

## Results

### Patients’ demographic and clinical characteristics

During the study period, a total of 333 cancer patients treated at least once in the department of medical oncology in Asan Medical Center (Seoul, Korea) responded to survey. The median age was 59 years (range, 29 to 70), and approximately half (51.7%) lived in rural areas more than 2 hours apart from the hospital. Colorectal cancer (35.4%) was the most common type of cancer, followed by stomach cancer (15.6%) and sarcoma (11.4%). Rare cancers constituted about 30% of the cohort, and sarcoma made the largest portion of rare cancers group. Most patients experienced metastatic disease (73.9%), and about half were initially diagnosed with metastatic cancer. While 64.9% had experience of relapse or disease progression, 23.7% of patients presented with a curable stage of disease without relapse. Nearly all patients (97.0%) received chemotherapy at least once, and more than half of the patients received at least two lines of chemotherapy. More than 70% of the patients in the cohort received surgical treatment regardless of the aim, which included 32.7% of palliative surgery.

In the comparison of common and rare cancers, significantly more of patients with common cancers experienced metastatic disease. Accordingly, common cancer group more frequently exposed to chemotherapy including two or more lines of treatment. Incidence of prior enrollment in CTs was also significantly higher in common cancer group (21.8% vs 12.4%, p=0.047) (Table 1).

**Table 1.** Patient characteristics

| **Characteristics** | **Total (%)** | **Common cancer**<br>**(n, %)** | **Rare cancer**<br>**(n, %)** | **P** |
| --- | --- | --- | --- | --- |
| Total patients | 333 (100) | 236 (70.9) | 97 (29.1) |  |
| **Age (median, range)** | 59 (2970) | 59 (29-70) | 58 (45-70) | 0.936 |
| **Gender** |  |  | 8 (66.7) | 0.011 |
| Male | 210 | 159 (67.4) | 51 (52.6) |  |
| Female | 123 | 77 (32.6) | 46 (47.4) |  |
| **Education degree** |  |  |  | 0.953 |
| Less than high school | 76 (23.0) | 55 (23.4) | 21 (21.9) |  |
| High school | 135 (40.8) | 95 (40.4) | 40 (41.7) |  |
| College or higher | 120 (36.3) | 85 (36.2) | 35 (36.5) |  |
| **Economic status <sup>a)</sup>** |  |  |  | 0.839 |
| < 18 | 84 (25.5) | 60 (25.8) | 24 (24.7) |  |
| ≥ 18 to < 45 | 145 (43.9) | 100 (42.9) | 45 (46.4) |  |
| ≥ 45 | 101 (30.6) | 73 (31.3) | 28 (28.9) |  |
| **Marital status <sup>b)</sup>** |  |  |  | 0.066 |
| Currently married | 47 (14.1) | 28 (11.9) | 19 (19.6) |  |
| Currently unmarried | 286 (85.9) | 208 (88.1) | 78 (80.4) |  |
| **Religion** |  |  |  | 0.989 |
| Christian | 86 (26.1) | 61 (26.1) | 25 (26.3) |  |
| Buddhist | 71 (21.6) | 51 (21.8) | 20 (21.1) |  |
| None/Others | 172 (52.3) | 122 (52.1) | 50 (52.6) |  |
| **Distance from clinic** |  |  |  | 0.384 |
| < 1hour | 78 (23.6) | 60 (25.4) | 18 (18.9) |  |
| ≥ 1hour to < 2hours | 82 (24.8) | 59 (25.0) | 23 (24.2) |  |
| ≥ 2 hours | 171 (51.7) | 117 (49.6) | 54 (56.8) |  |
| **Cancer type** |  |  |  | <0.001 |
| Colorectal | 118 (35.4) | 118 (50.0) | 0 (0.0) |  |
| Gastric | 52 (15.6) | 52 (22.0) | 0 (0.0) |  |
| Lung | 35 (10.5) | 35 (14.8) | 0 (0.0) |  |
| Breast | 2 (0.6) | 2 (0.8) | 0 (0.0) |  |
| Pancreatobiliary | 19 (5.7) | 14 (5.9) | <sup>c)</sup>5 (5.2) |  |
| Genitourinary | 17 (5.1) | 15 (6.4) | <sup>d)</sup>2 (2.1) |  |
| Sarcoma | 38 (11.4) | 0 (0.0) | 38 (39.2) |  |
| Neuroendocrine tumor | 19 (5.7) | 0 (0.0) | 19 (19.6) |  |
| Other rare tumors | 33 (9.9) | 0 (0.0) | 33 (34.0) |  |
| **Disease status** |  |  |  | <0.001 |
| Initially metastatic | 174 (52.3) | 136 (57.6) | 38 (39.2) |  |
| Relapsed metastatic | 72 (21.6) | 48 (20.3) | 24 (24.7) |  |
| Curable | 87 (26.1) | 52 (22.0) | 35 (36.1) |  |
| **Experience of metastasis** |  |  |  | 0.008 |
| No | 87 (26.1) | 52 (22.0) | 35 (36.1) |  |
| Yes | 246 (73.9) | 184 (78.0) | 62 (63.9) |  |
| **Relapse or progression** |  |  |  | 0.214 |
| No | 117 (35.1) | 100 (42.4) | 59 (60.8) |  |
| Yes | 216 (64.9) | 136 (57.6) | 38 (39.2) |  |
| **Experience of CTx <sup>e)</sup>** |  |  |  | <0.001 |
| No | 10 (3.0) | 1 (0.4) | 9 (9.3) |  |
| Yes | 323 (97.0) | 235 (99.6) | 88 (90.7) |  |
| **Median number of CTx lines (range)** | 2 (0-7) | 2 (0-7) | 1 (0-6) | 0.002 |
| **\*CTx lines ≥ 2** |  |  |  | 0.004 |
| No | 151 (45.4) | 95 (40.3) | 56(57.7) |  |
| Yes | 182 (54.7) | 141 (59.7) | 41(42.3) |  |
| **Surgical treatment** |  |  |  | 0.264 |
| No | 86 (25.8) | 65 (27.5) | 21 (21.6) |  |
| Yes | 247 (74.2) | 171(72.5) | 76 (78.4) |  |
| **\*Aim of surgery** |  |  |  | 0.184 |
| Curative | 165 (67.4) | 110 (64.7) | 55 (73.3) |  |
| Palliative | 80 (32.7) | 60 (35.3) | 20 26.7) |  |
| **Experience of CTs enrollment** |  |  |  | 0.047 |
| No | 268 (81.0) | 183 (78.2) | 85 (87.6) |  |
| Yes | 63 (19.0) | 51 (21.8) | 12 (12.4) |  |

CTx; chemotherapy, CTs; clinical trials
<sup>a)</sup> Units are 100 US dollars per month <sup>b)</sup> Currently unmarried state includes both single and divorced state <sup>c)</sup> Cholangiocarcinoma, <sup>d)</sup> Upper tract urothelial carcinoma, <sup>e)</sup> Includes all types of chemotherapy regardless of the aim or disease setting

### Current perceptional status and influential factors of CTs in patients with common and rare cancer

In the cohort, 87.7% and 75.3% of respondents answered they had heard of and knew about CTs, respectively, and 55.6% expressed willingness to participate in CTs. These perceptional parameters were significantly inter-correlated (p <0.001, data not shown), and were consistently related to prior enrollment in CTs. In particular, patients’ willingness was even more strongly influenced by awareness and perception, as well as their prior enrollment experiences (Table S1). Patients with common cancers showed significantly higher levels of awareness and perception (79.9% vs 64.2%, p=0.003 and 91.9% vs 77.3%, p<0.001), as well as willingness to participate in CTs (58.9% vs 47.4%, p=0.06) than those with rare cancers (Table 2). Regardless of generally favorable perceptions, the majority was not heard of or knew about ongoing CTs for their type of cancer, which was more obvious in rare cancer group (p=0.006). Most patients also did not know other patients with experience of participation in CTs, and only about a third of the patients had experience of being encouraged or advised to participate in CTs.

**Table 2.** Comparison of key perceptional parameters related to clinical trials (CTs) between patients with common and rare cancers

| **Perceptual outcomes** | **Total** | **Common cancers** | **Rare cancers** | ***p*** |
| --- | --- | --- | --- | --- |
| **Heard about CTs *in general*** | 292 (87.7) | 217 (91.9) | 75 (77.3) | <0.001 |
| **Heard about CTs for *your disease*** | 123 (37.2) | 98 (41.9) | 25 (25.8) | 0.006 |
| **Heard advice to participate in CTs from anyone** | 106 (31.8) | 82 (34.7) | 24 (24.7) | 0.075 |
| **Knew about CTs** | 244 (75.3) | 183 (79.9) | 61 (64.2) | 0.003 |
| **Knew anyone who participated in CTs** | 60 (18.1) | 43 (18.3) | 17 (17.5) | 0.868 |
| **Made efforts to collect information about your disease** | 31 (9.3) | 23 (9.7) | 8 (8.2) | 0.669 |
| **Expressed willingness to participate in CTs** | 185 (55.6) | 139 (58.9) | 46 (47.4) | 0.056 |

Except for the cancer type, perception and awareness were not affected by most of the variables in both common and rare cancer groups. However, willingness was significantly influenced by disease status and experience of metastasis (p<0.01), and higher education level also showed statistical correlation with willingness (p=0.05). (Table 3)

**Table 3.**
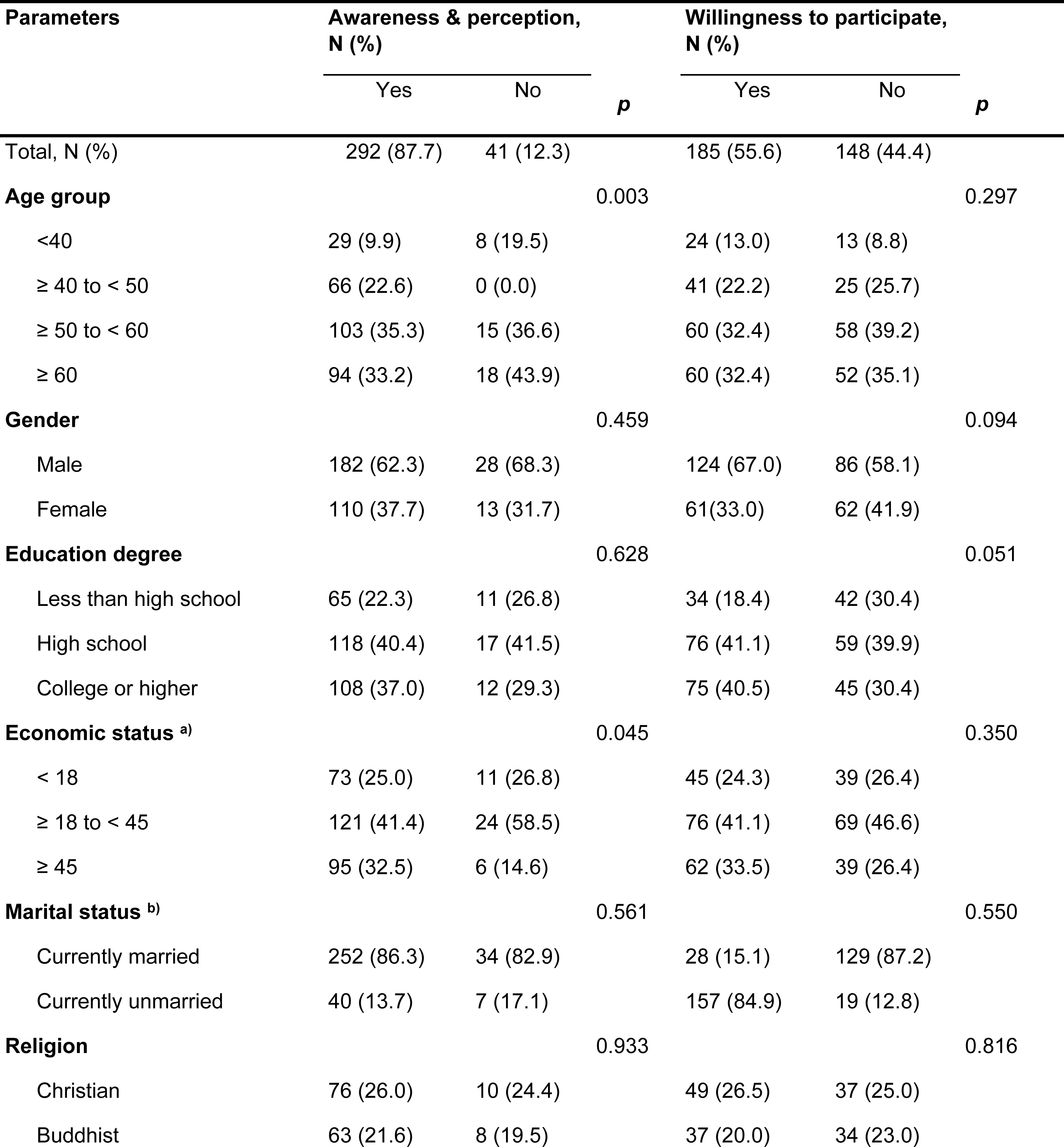
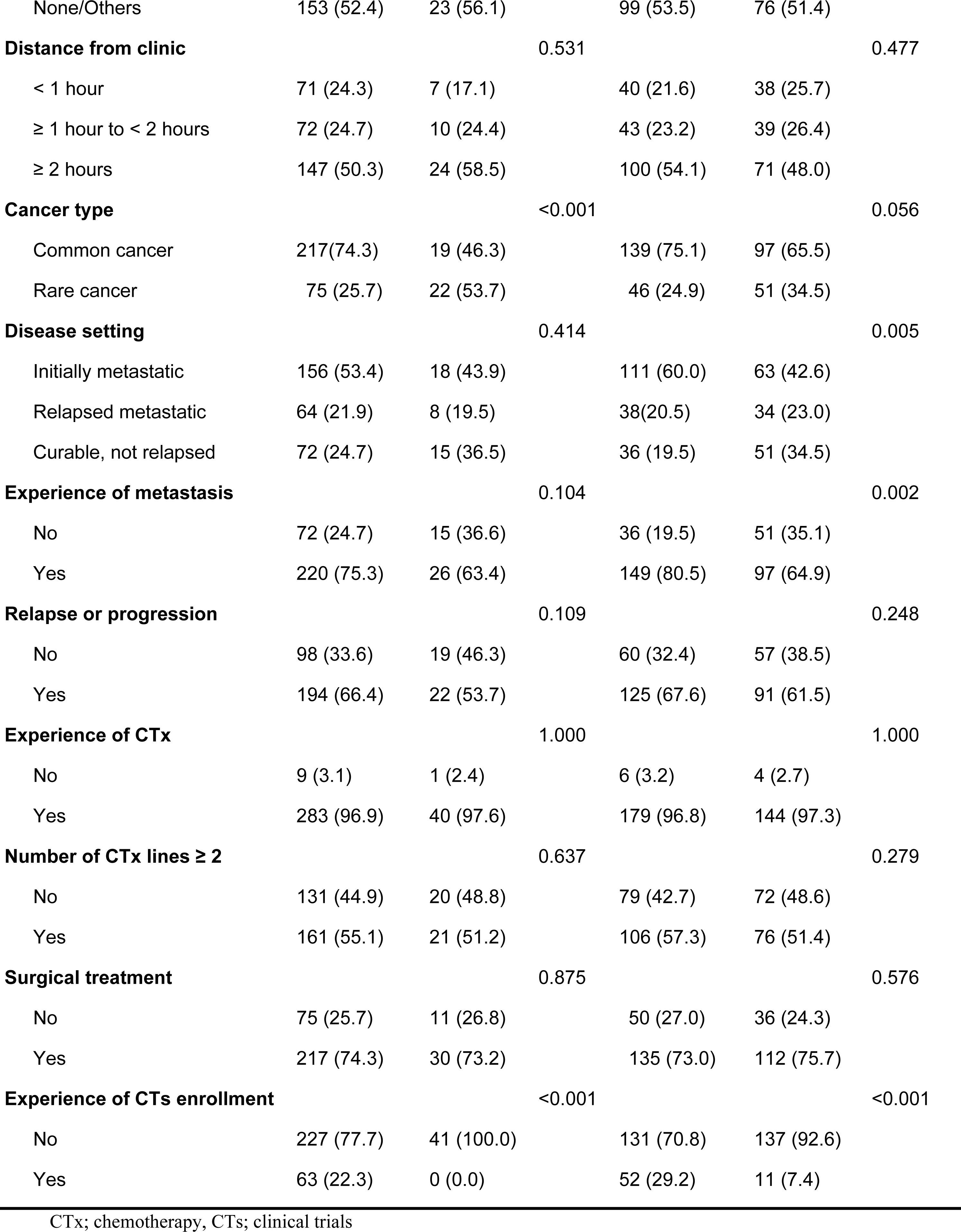
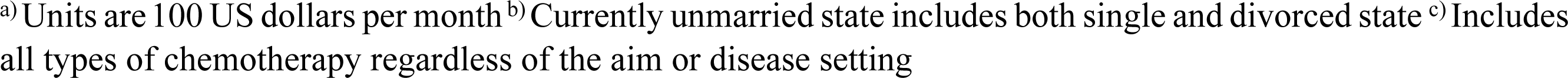
Influential clinical and demographic parameters of cancer patients’ awareness and perception of clinical trials and willingness to participate in them

### Comparison of perceptional parameters between patients with common and rare cancers using the PARTAKE survey

A comparative analysis was performed to further evaluate whether there are perceptional differences regarding CTs between common and rare cancer groups using the PARTAKE survey (Table 4).

**Table 4.** Comparison of perceptional structures regarding clinical trials (CTs) between patients with common and rare cancers

| No. | Questionnaire | Common cancer<br>(N=236) | Rare cancer<br>(N=97) | Total<br>(N=333) | p |
| --- | --- | --- | --- | --- | --- |
| **Value : perceptions regarding the value that research brings** |  |  |  |  |  |
| V1 | Clinical research benefits society |  |  |  | 0.72 |
|  | True | 196 (83.1) | 78 (80.4) | 274 (82.3) |  |
|  | False | 8 (3.4) | 5 (5.2) | 13 (3.9) |  |
|  | Not aware | 32 (13.5) | 14 (14.4) | 46 (13.8) |  |
| V2 | Clinical research harms society |  |  |  | 0.27 |
|  | True | 11 (4.7) | 7 (7.2) | 18 (5.4) |  |
|  | False | 202 (85.6) | 76 (78.4) | 278 (83.5) |  |
|  | Not aware | 23 (9.7) | 14 (14.4) | 37 (11.1) |  |
| V3 | Clinical research is an essential step in developing new treatment |  |  |  | < 0.01 |
|  | True | 224 (94.1) | 81 (83.5) | 303 (91.0) |  |
|  | False | 1 (0.4) | 1 (1.0) | 2 (0.6) |  |
|  | Not aware | 13 (5.5) | 15 (15.5) | 28 (8.4) |  |
| V4 | Hospitals that participate in clinical research provide better healthcare |  |  |  | 0.69 |
|  | True | 125 (53.0) | 48 (49.5) | 173 (52.0) |  |
|  | False | 14 (5.9) | 8 (8.2) | 22 (6.6) |  |
|  | Not aware | 97 (41.1) | 41 (42.3) | 138 (41.4) |  |
| V5 | Experiments on humans are essential to developing new treatments |  |  |  | < 0.01 |
|  | True | 185 (78.4) | 61 (62.9) | 246 (73.9) |  |
|  | False | 15 (6.4) | 6 (6.2) | 21 (6.3) |  |
|  | Not aware | 36 (15.3) | 30 (30.9) | 66 (19.8) |  |
| **Motivation : perceptions regarding reasons for doing or participating in research** |  |  |  |  |  |
| M1 | The most important reason for developing new treatments is the advancement of science |  |  |  | 0.12 |
|  | True | 196 (83.1) | 73 (77.3) | 271 (81.4) |  |
|  | False | 15 (6.4) | 4 (4.1) | 19 (5.7) |  |
|  | Not aware | 25 (10.6) | 18 (18.6) | 43 (12.9) |  |
| M2 | The most important reason for developing new treatments is financial gain |  |  |  | 0.18 |
|  | True | 63 (26.8) | 33 (34.0) | 96 (28.9) |  |
|  | False | 125 (53.2) | 52 (53.6) | 177 (53.3) |  |
|  | Not aware | 47 (20.0) | 12 (12.4) | 59 (17.8) |  |
| M3 | Participation in research is entirely voluntary |  |  |  | 0.35 |
|  | True | 160 (67.8) | 58 (59.8) | 218 (65.5) |  |
|  | False | 30 (12.7) | 14 (14.4) | 44 (13.2) |  |
|  | Not aware | 46 (19.5) | 25 (25.8) | 71 (21.3) |  |
| M4 | Altruism is the only valid reason for participation in research |  |  |  | 0.42 |
|  | True | 59 (25.1) | 18 (18.6) | 77 (23.2) |  |
|  | False | 101 (43.0) | 44 (45.4) | 145 (43.7) |  |
|  | Not aware | 75 (31.9) | 35 (36.1) | 110 (33.1) |  |
| M5 | You have had an opportunity to participate in clinical research |  |  |  | 0.46 |
|  | True | 77 (32.6) | 25 (25.8) | 102 (30.6) |  |
|  | False | 138 (58.5) | 62 (63.9) | 200 (60.1) |  |
|  | Not aware | 21 (8.9) | 10 (10.3) | 31 (9.3) |  |
| **Compliance : perceptions regarding the conduct of research** |  |  |  |  |  |
| C1 | Volunteers in clinical research get adequate compensation for their participation |  |  |  | 0.38 |
|  | True | 44 (18.6) | 18 (18.6) | 62 (18.6) |  |
|  | False | 26 (11.0) | 16 (16.5) | 42 (12.6) |  |
|  | Not aware | 166 (70.3) | 63 (64.9) | 229 (68.8) |  |
| C2 | Participants in clinical research get adequate compensation for any adverse outcomes |  |  |  | 0.85 |
|  | True | 47 (19.9) | 22 (22.7) | 69 (20.7) |  |
|  | False | 34 (14.4) | 14 (14.4) | 48 (14.4) |  |
|  | Not aware | 155 (65.7) | 61 (62.9) | 216 (64.9) |  |
| C3 | Confidentiality of research participants is adequately protected |  |  |  | 0.08 |
|  | True | 134 (57.5) | 43 (44.3) | 177 (53.6) |  |
|  | False | 12 (5.2) | 5 (5.2) | 17 (5.2) |  |
|  | Not aware | 87 (37.3) | 49 (50.5) | 136 (41.2) |  |
| C4 | Volunteers in clinical research get adequate information about the research they participate in |  |  |  | 0.98 |
|  | True | 106 (44.9) | 43 (44.3) | 149 (44.7) |  |
|  | False | 25 (10.6) | 11 (11.3) | 36 (10.8) |  |
|  | Not aware | 105 (44.5) | 43 (44.3) | 148 (44.4) |  |
| C5 | Researchers make sure research is safe for participants |  |  |  | 0.22 |
|  | True | 124 (52.8) | 43 (44.3) | 167 (50.3) |  |
|  | False | 9 (3.8) | 7 (7.2) | 16 (4.8) |  |
|  | Not aware | 102 (43.4) | 47 (48.5) | 149 (44.9) |  |
| C6 | Harmful events occurring during a clinical trial must be due to experimental treatment |  |  |  | 0.63 |
|  | True | 91 (38.6) | 32 (33.0) | 123 (36.9) |  |
|  | False | 28 (11.9) | 12 (12.4) | 40 (12.0) |  |
|  | Not aware | 117 (49.6) | 53 (54.6) | 170 (51.1) |  |
| C7 | The public should be involved in clinical research (e.g., design, oversight, funding) |  |  |  | 0.22 |
|  | True | 80 (34.0) | 24 (24.7) | 104 (31.3) |  |
|  | False | 48 (20.4) | 25 (25.8) | 73 (22.0) |  |
|  | Not aware | 107 (45.5) | 48 (49.5) | 155 (46.7) |  |
| **Trust : perceptions regarding trust of research entities** |  |  |  |  |  |
| T1 | The government always adequately protects the public against unethical clinical research |  |  |  | 0.01 |
|  | True | 78 (33.1) | 16 (16.5) | 94 (28.2) |  |
|  | False | 33 (14.0) | 17 (17.5) | 50 (15.0) |  |
|  | Not aware | 125 (53.0) | 64 (66.0) | 189 (56.8) |  |
| T2 | Clinical research information provided by pharmaceutical companies can be trusted |  |  |  | 0.06 |
|  | True | 84 (35.6) | 30 (30.9) | 114 (34.2) |  |
|  | False | 31 (13.1) | 23 (23.7) | 54 (16.2) |  |
|  | Not aware | 121 (51.3) | 44 (45.4) | 165 (49.5) |  |
| T3 | Clinical research information provided by academic institutions can be trusted |  |  |  | <0.01 |
|  | True | 138 (58.5) | 48 (50.0) | 186 (56.0) |  |
|  | False | 7 (3.0) | 12 (12.5) | 19 (5.7) |  |
|  | Not aware | 91 (38.6) | 36 (37.5) | 127 (38.3) |  |
| T4 | If you decide not to participate in research, your doctor will not give you good care |  |  |  | 0.99 |
|  | True | 16 (6.8) | 7 (7.3) | 23 (6.9) |  |
|  | False | 170 (72.0) | 69 (71.9) | 239 (72.0) |  |
|  | Not aware | 50 (21.2) | 20 (20.8) | 70 (21.1) |  |
| T5 | Doctors force their patients to participate in research |  |  |  | 0.58 |
|  | True | 8 (3.4) | 3 (3.1) | 11 (3.3) |  |
|  | False | 184 (78.0) | 71 (73.2) | 255 (76.6) |  |
|  | Not aware | 44 (18.6) | 23 (23.7) | 67 (20.1) |  |
| T6 | Human participants in clinical research are treated like experimental animals ('human guinea pigs') |  |  |  | 0.38 |
|  | True | 18 (7.6) | 11 (11.3) | 29 (8.7) |  |
|  | False | 130 (55.1) | 56 (57.7) | 186 (55.9) |  |
|  | Not aware | 88 (37.3) | 30 (30.9) | 118 (35.4) |  |
| T7 | Confidentiality is a matter of importance to research participants |  |  |  | 0.81 |
|  | True | 158 (66.9) | 67 (69.1) | 225 (67.6) |  |
|  | False | 53 (22.5) | 22 (22.7) | 75 (22.5) |  |
|  | Not aware | 25 (10.6) | 8 (8.2) | 33 (9.9) |  |
| T8 | All the results of clinical research are made available to the public |  |  |  | 0.55 |
|  | True | 32 (13.6) | 9 (9.3) | 41 (12.3) |  |
|  | False | 95 (40.3) | 40 (41.2) | 135 (40.5) |  |
|  | Not aware | 109 (46.2) | 48 (49.5) | 157 (47.1) |  |
| T9 | The media accurately describes clinical research |  |  |  | 0.16 |
|  | True | 62 (26.3) | 16 (16.5) | 78 (23.4) |  |
|  | False | 53 (22.5) | 24 (24.7) | 77 (23.1) |  |
|  | Not aware | 121 (51.3) | 57 (58.8) | 178 (53.5) |  |

#### Value and motivation

Most respondents agreed about the social and scientific benefits of CTs but it was better perceived by patients with common cancers (V1, V3, V5). The most important reason for participation was personal medical benefits from CTs (68.8%), followed by trust in investigators (33.3%) in both common and rare cancer groups. In contrast, concern about safety (61.4%) and fear of CTs (31.0%) were the two most concordant reasons for refusal. Notably, 40% of patients with rare cancers do not believe or do not know if enrollment in CTs is entirely voluntary, and 14.8% and 14.5% of patients answered that they would refuse CTs because of the lack of enrollment opportunities and sufficient knowledge, respectively. Although there was no demonstrable difference between patients with common and rare cancers regarding these reasons, patients with rare cancers were more frequently motivated by altruism, and negatively affected by fear and ignorance than the patients with common cancers (Figure 1).

**Figure 1.**
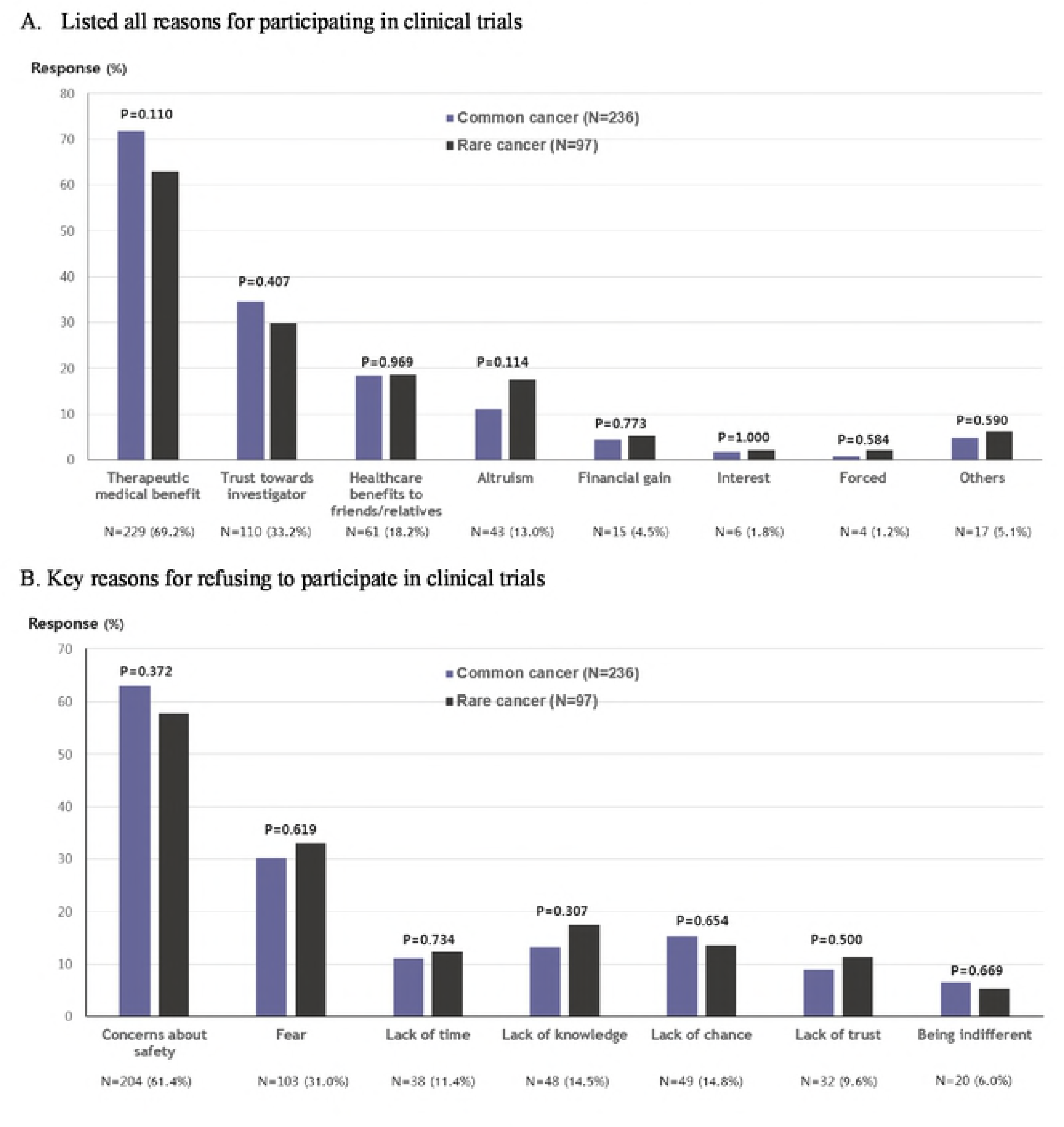
Comparison or reasons Cor participating in and refusing clinical trials between patients with common arid rare cancers (*allowed multiple choices)

#### Compliance and Trust

About half of the patients answered that the researchers would guarantee safety (C5) and provide adequate information during the conduct of CTs (C4). However, the majority did not have proper concept of reward from their participation in CTs and adverse events (C1, C2). Medical doctors were most trusted sources of credible information related to CT in both common and rare cancer groups. Meanwhile, pharmaceutical companies and the media were significantly more trusted by patients in the common cancers group, whereas government was more trusted by the rare cancer patients (Figure 2). Patients were highly dependent on their investigators for deciding enrollment in CTs, but they generally did not consider their physicians as compelling them to participate in CTs (T5), and most respondents trusted that doctors would not exclude them from an optimal treatment opportunity even after the refusal of CTs (T4).

#### Comparison of perceptional parameters in cancer patients and the general public

Comparing with our previous study in the Korean general public(5), perceptional parameters were significantly more favorable in cancer patients than in the general public (Table S2). Cancer patients better acknowledged the medical, scientific and social needs of CTs in terms of newer drug development than the general public, while the public exhibited greater concerns about the issues of safety and confidentiality, as well as the information and rewards provided (Table S3). However, clinical benefit was the most important reason for enrollment in CTs in both groups. Meanwhile, cancer patients were more influenced by trust in their investigator (33.2% vs 18.0%, p<0.001), whereas the general public was more influenced by private financial and intellectual interests from CTs (31.8% vs 4.5%, p<0.001) (Figure S1).

## Discussion

Present study depicts the current favorable perceptional status regarding CTs in Korean cancer patients with high levels of awareness and perception. However, willingness of participating in CTs was only comparable rather than favorable although it was significantly correlated with awareness and perception. Willingness was also strongly correlated with previous experience of enrollment, more obviously in rare cancer group.

The salience of our investigation lies in further comparison of these perceptional parameters between the patients with common and rare cancers, which surprisingly reveals generally less positive perception and willingness of CTs in patients with rare cancer patients. Comparative analysis of the results of the PARTAKE survey additionally alludes a tendency of more widespread ignorance, mistrust and hesitance with fear of CTs in rare cancer group. In addition, substantial patients of rare cancer group exhibited indifference or mistrust about voluntariness of participation in CTs, which might hinder their access to CTs. This gives a sobering message of a challenge that we should make further enrichment of willingness as well as perception of CTs in patients with rare cancers. They need to be properly educated and encouraged to feel safe and confident of CTs, and then should be exposed to expanded opportunities for enrollment.

Our study suggests a substantial gap between perception and knowledge about CTs in cancer patients despite generally improved perceptional status. To our surprise, more than half of the cancer patients were not aware of currently ongoing CTs for their own diseases, and this was more prominent in rare cancer patients. Accordingly, nearly 15% of all cancer patients would refuse CTs for the lack of adequate information or opportunity of enrollment. Considering that cancer patients were greatly influenced by their investigators and the information they provide when deciding enrollment, investigator’s efforts for informing and motivating their patients to participate in CTs could primarily contribute to narrowing this gap.

There is an inevitable methodological limitation of a single center study. However, perceptional outcomes in our cohort generally accorded with other similar studies (2-4, 6, 11, 23, 24). Also, given that our institute places one of the largest tertiary cancer centers in Korea with a high volume of CTs, and that it is the latest study investigating perceptional aspects of CTs in cancer patients, the study might promptly provide updated insights on CTs. Accordingly, we observed incremental progress of perceptional status toward CTs in cancer patients compared with several previous Korean and other studies (1-4).

Current study arouses timely focus on the rare cancer population, which might be another illuminating part of the study. Newly developed CTs designs based on the advancement of sequencing technologies have broadened opportunities of CTs in patients with rare cancers(19, 25). Accordingly, numbers of currently registered and ongoing CTs for rare cancers on ClinicalTrials.gov have decently increased compared with last three decades (Table S4) (14, 20, 26). However, we do not exactly know what make patients with rare cancers participating in CTs and what make them refraining from it. Relatively unfavorable perceptional status with less willingness observed in the study should be reminded by research entities to construct more successful strategy of CTs in patients with rare cancers. Furthermore, organized efforts for rare cancers should be steadily continued to overcome perceptional obstacles lying in rare cancers (27-29).

In summary, our study demonstrates recent perceptional advancement but also illuminates current unmet need of CTs in cancer patients by revealing relative perceptional poverties in patients with rare cancers. Perception may possibly shape behavior(5), but it could not alone guarantee the success of CTs in cancer patients, especially for the rare cancers. Therefore, in order to convert patients’ perceptions to genuine voluntariness in the era of CTs, it seems necessary to enrich knowledge and willingness of CTs through further education and encouragement by research and public entities, which might be more critical to populations suffering rare cancers. Finally, we should consistently pursue the optimal chances of CTs enrollment for benefiting patients with both rare and common cancers. This would lessen long-lasted therapeutic gap between common and rare cancers, and we might eventually step forward to a newer trajectory of cancer treatment.

## Acknowledgment

The current research was supported by two grants from the Korea Health Technology R&D Project of the Korea Health Industry Development Institute (KHIDI), funded by the Ministry of Health & Welfare, Republic of Korea (grant number: HI14C1061, HI18C2383)

